# Inverse Density of Iba1 and Glutamine Synthetase Expressing Glia in Rat Inferior Colliculus

**DOI:** 10.1101/2021.06.30.450495

**Authors:** Llwyd David Orton

## Abstract

Microglia and astrocytes undertake numerous essential roles in nervous systems but we know little of their anatomical distribution within numerous nuclei. In the principal nuclei of the mammalian auditory midbrain, the inferior colliculi (IC), the cellular density and relative distribution of glutamate synthetase (GS) expressing astrocytes and ionized calcium-binding adapter molecule 1 (Iba1) expressing microglia is unknown. To address this, the IC of young adult, male Wistar rats were immunohistochemically labelled for GS and Iba1, using chromogenic methods. Sub-regions of imaged IC sections were demarked and soma density of both cell types determined. GS labelled somata were twice more densely packed as Iba1 labelled somata throughout IC parenchyma and peri-vascular regions. Furthermore, GS labelled somata density was significantly lower in dorsal cortex than external cortex or central nucleus. Iba1 labelled somata density exhibited the opposite trend, revealing an inverse density of these glial cell types between IC sub-regions. GS labelled neuropil was strongest in the cortices with and a gradual transition of lighter labelling towards central nucleus. These data provide the first detailed descriptions of GS labelling in IC and demonstrate sub-regional differences in IC glial cell density. Taken together, these findings suggest neurochemical specialization of glia in IC sub-regions, likely related to local physiological and metabolic demands, with implications for IC function.

## Introduction

The inferior colliculi (IC) are the principal nuclei of the sub-cortical auditory pathway. Cellular processing in IC integrates afferent drive from a diversity of ascending (Adams, 1979), descending (Bajo et al., 2007; Patel et al., 2017) and commissural sources (Malmierca et al., 2009; Orton et al., 2016, 2012). The IC contains a laminar arrangement of cells and connections not replicated elsewhere in the brain, necessitating their detailed characterisation. While the overwhelming majority of our knowledge about the IC relates to neuronal anatomy, physiology and connectivity, glia comprise a significant proportion of cells in the central nervous system (Herculano-Houzel, 2014) and our understanding of glia in IC lags far behind other brain regions. In particular, astrocytes and microglia in IC remain largely uncharacterised.

Astrocytes perform a multitude of roles during ongoing processing and may express a number of somatic protein markers. It has been shown that in young IC of healthy gerbil (Hafidi and Galifianakis, 2003) and guinea pig (Webb and Orton, 2020), the most widely used astrocyte antibody, glial fibrillary acidic protein (GFAP) is expressed strongly in the *glia limitans externa* and the outermost layers of IC, with few cells labelled in the parenchyma. Sulforhodamine 101 (SR101), which preferentially identifies astrocytes, has recently been shown to label parenchymal somata in IC (Ghirardini et al., 2018), as does S100 (Hafidi and Galifianakis, 2003), but the distribution of other markers is unknown.

Ramified microglia tile the brain parenchyma during ongoing processing with heterogeneous morphologies (Lawson et al., 1990; Olah et al., 2011; Tan et al., 2020). The population of ionized calcium-binding adapter molecule 1 (Iba1) labelled microglia vary in the extent of ramifications between IC sub-regions in guinea pig (Webb and Orton, 2020) and are activated in response to contralateral deafening (Rosskothen-Kuhl et al., 2018).

Recent findings in the somatosensory (Kwak et al., 2020; Lines et al., 2020) and visual (Cheadle et al., 2020; Tremblay et al., 2010) pathways have shown the essential, active roles glia play in sculpting ongoing neural processing of sensory stimuli. In the auditory pathway, electrical stimulation of cochlea induces activation of both astrocytes and microglia in IC (Rosskothen-Kuhl et al., 2018), suggesting these glia also play essential roles in IC processing. Consequently, there is a need to characterise the anatomical distribution of glial sub-types. To the author’s knowledge, there have been no detailed descriptions of GS astrocytes in IC and furthermore, the relative cellular density of microglia and astrocytes is unknown. Therefore, this study sought to characterise the somatic and neuropil labelling density of two glial sub-types: glutamine synthetase (GS) expressing astrocytes and Iba1 expressing microglia in the young adult rat brain.

## Materials and methods

### Regulation and Ethics

Ethical approval was provided by the Faculty Research Ethics and Governance Committee in the Faculty of Science and Engineering at Manchester Metropolitan University (EthOS reference numbers: 401 & 12,561). Seven male Wistar rats were used, weighing 150-199g, aged 8-10 weeks. Animals were euthanised via cervical dislocation in line with the Animals (Scientific Procedures) Act 1986 by an appropriately trained person. The number of animals used and their suffering was minimised.

### Tissue processing

Brains were removed with rongeurs (Micro Friedman, 0.8mm jaws, WPI) and a scalpel and placed in 4% paraformaldehyde in 0.1M phosphate buffered saline (PBS) at 4°C for at least one month, to ensure thorough fixation. Brains were transected in the coronal plane, rostral to the superior colliculus, and placed in 30% sucrose in PBS at 4°C, until they sank. The tissue block was then placed in an embedding mold (Peel-a-way; Shandon), covered in embedding medium (OCT; Agar Scientific) and frozen at −80°C until sectioning.

Each brain block was sectioned serially in the coronal plane at 50μm on a cryostat (Leica CM 3050 S) in a rostro-caudal direction throughout the extent of the IC. The starting point of the first section was randomly selected (Gundersen and Jensen, 1987). Serial sections were collected in a PBS filled 12-well plate such that every well contained a series of sections separated by 600μm. Every 6^th^ section through the IC was sampled in a systematic-uniform-random manner with the first well selected at random. This procedure produced 7-10 sections through each IC, spaced at 300μm intervals. The total sampling fraction of each IC was therefore one-sixth, in line with established best practice (Long et al., 1998). Accurate estimation of cell density can be obtained from this number of sections at this sampling fraction (West et al., 1991). This approach allowed for reconstruction of labelled sections in position order through each IC.

### Immunohistochemical labelling

The following primary antibodies were used:

Rabbit anti-Iba1 (1:1,000; polyclonal; 019-19741; Wako; RRID: AB_839504) - according to the manufacturer, this affinity purified antibody was raised against a synthetic peptide corresponding to the C-terminus fragment of rat Ionized calcium binding adaptor molecule 1 (Iba1). Labelling via western blot was positive for a 17kDa band (Imai et al., 1996). Selective labelling of ramified microglia was observed, matching similar reports rat (Fuentes-Santamaría et al., 2012) and guinea pig (Webb and Orton, 2020).

Mouse anti-Glutamine Synthetase (1:500; monoclonal; AB64613; Abcam; RRID: AB_1140869) – according to the manufacturer, the immunizing antigen is a recombinant fragment with tag: IEKLSKRHQY HIRAYDPKGG LDNARRLTGF HETSNINDFS AGVANRSASI RIPRTVGQEK KGYFEDRRPS ANCDPFSVTE ALIRTCLLNE TGDEPFQYKN, corresponding to amino acids 274□374 of human glutamine synthetase. Blotting detects a band of approximately 37 kDa (Heller et al., 2017). Previous labelling has shown this antibody is a selective marker of glial cells in rat (Huang et al., 2013).

All steps in the labelling protocol involved continuous gentle agitation of sections. Free-floating sections were washed 3×5mins in PBS. Sections were then incubated in 0.3% hydrogen peroxide in methylated spirits to block endogenous peroxidases. Following three more washes in PBS, sections were incubated in sodium citrate buffer (pH 6.0) for 20 minutes at 80°C in a water bath. After three more washes in PBS, sections were blocked in 5% normal goat serum (Vector) for one hour at room temperature. Subsequent to blocking, sections were incubated in Iba1 antibody in blocking solution, overnight at room temperature. The next day, sections were washed 3×5mins in PBS and incubated in biotinylated anti-rabbit IgG (1:1,000; Vector) in blocking solution for 1 hour. After 3xPBS washes, pre-mixed avidin-HRP (each 1:500) was applied for 30 minutes. After 3xPBS washes, Iba1+ labelling was produced via application of DAB (Vector). Sections were then taken through the same process for GS labelling with an antimouse secondary antibody and nickel intensified DAB. Sections were mounted on gelatin-chrome alum coated slides, dehydrated, cleared and coverslipped in DPX (Sigma). All sections included in this study were stained concurrently, with controls including exclusion of the primary, secondary and both antibodies on separate sections, which were free of cellular and neuropil labelling.

### Image acquisition

Sections were imaged on a 3D Histec Panoramic 250 slide scanner at 40x. All imaging parameters were the same for all images. Sub-regions of the IC on each section were annotated in CaseViewer software with reference to a rat brain atlas (Paxinos et al., 2009). Demarking the outer border of the IC and respective sub-regions throughout the reference volume enabled estimation of whole IC and sub-region volume via the Cavalieri method.

### Somata count analyses

Somata counts were performed within defined outer borders of each IC sub-region. Field of view locations were defined at positions within each IC sub-region on each section. Only sections where the entire counting frame was filled with parenchymal tissue were counted. Cell counts were only performed on parenchymal regions where no vessels were present. The x-y dimensions of the field of view for Iba1 cell counts was 100μm^2^ and for GS it was 25μm^2^. Based on pilot data, it was calculated that to count a minimum of 100 cells per animal required two fields of view per sub-region per section for Iba1+ somata and one for GS+ somata. To allow comparisons of respective cell densities, GS+ soma counts were multiplied by a factor of 4 to produce comparable estimates of cell density per 100μm^2^. Cells were subject to standard counting rules - i.e. only counted if some part of the soma was within the field of view but not if any part of the soma touched the left or lower borders. The most in-focus point of each cell was only counted once to remove double counting errors. If field of view placement produced adjacent probe frames, the exclusion lines were extended across the entire IC such that cells were only counted once. For region of interest analyses, cell locations were demarked using the *Multi-Point* tool in ImageJ, from which 2D Delaunay triangulations and Voronoi tessellations were derived.

For peri-vascular analyses, one longitudinal blood vessel was located in each sub-region of each section. The 2D length of the vessel was measured using the *Linear Measurement* tool in CaseViewer. All labelled peri-vascular somata were identified and the 2D median distance between somata was calculated for that vessel. A small percentage of longer vessels curved such that the linear measurement tool underestimated their length, so the *Open Polygon* tool was used. Somata counts were then applied along the vessel length following the above approach, with the addition requirement that each counted soma must directly abut the vessel.

### Neuropil density analyses

Whole IC images (5x) and regions of interest from each IC sub-region (40x) were exported as tiff images and imported into ImageJ. Whole IC images from each animal were selected as being midway along the rostro-caudal axis and were converted to 8-bit format. The *Straight Line* tool was placed along two axes: 1) along the dorso-ventral axis of the IC, from the *glia limitans externa* to the ventral border of the IC to the underlying lateral lemniscus; and 2) along the medio-lateral axis, from the *glia limitans externa* to the medial border with the periaqueductal grey. The *Plot Profile* analysis quantified grey levels for each pixel along each axis. Three parallel lines were placed along each axis for each IC image and the mean was calculated.

### Statistical analyses

Data were collected in LibreOffice. Graphs were generated and statistical hypothesis testing was performed in Prism (GraphPad). A two-way ANOVA was used to test for interactions between cell type (factor 1) and distance from the caudal pole (factor 2) with comparisons between cell means and every other cell mean and a standard α criterion of 0.05. Multiple comparison tests were performed using the Tukey method. Welch’s one-way ANOVAs were conducted for a number of single factor analyses with Dunnett T3 multiple *post-hoc* comparisons, where appropriate. P values were multiplicity adjusted for *post-hoc* analyses. Correlations were performed using the Pearson method.

## Results

Both antibodies consistently labelled distinct somata and neuropil, with Iba1 additionally labelling ramified processes of putative microglia. Neuropil labelling for Iba1+ was of low intensity and similar throughout the tissue section. Conversely, GS+ neuropil labelling exhibited marked differences between IC sub-regions. Figure 1 shows four serial low power (5x) micrographs of coronal sections from animal RR4, showing characteristic labelling. The borders of IC sub-regions were delineated with reference to the atlas of Paxinos et al., (2009) which sub-divided the IC into dorsal cortex (DC), central nucleus (CN) and external cortex (EC). At low resolution, the differences observed between sub-regions in GS+ neuropil labelling are apparent at the rostral-most section (Fig. 1A). At this position through the pons, strong labelling was also found in the superficial layers of the superior colliculus. More moderate labelling was found in the peri-aqueductal grey, deep layers of superior colliculus and brachium of the IC. Lighter labelling was found in rostral EC. Also apparent was the strong Iba1+ cellular labelling in microglia forming the *glia limitans externa* and the peri-aqueductal border.

**Figure 1.**
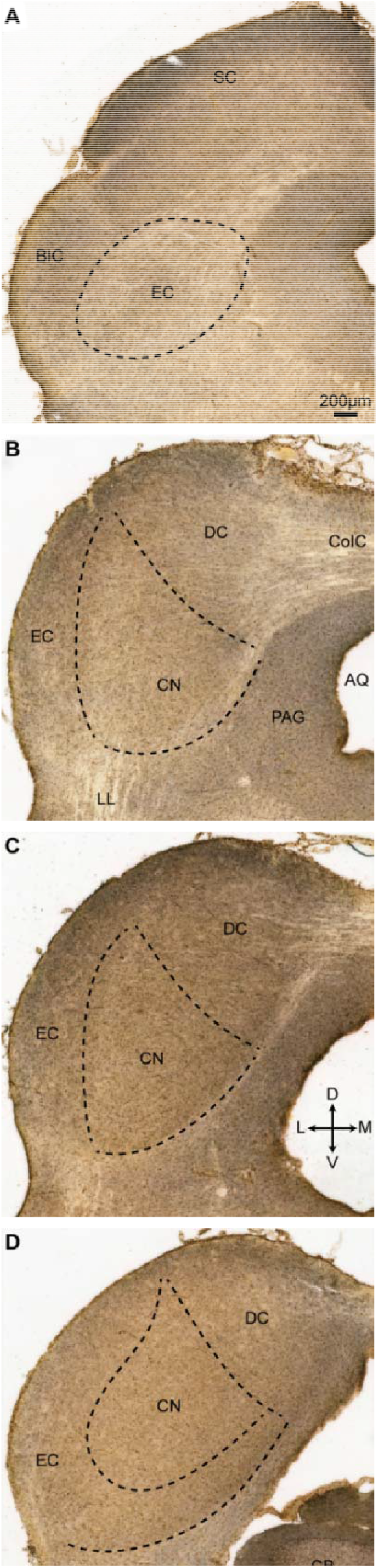
Low power (5x) photomicrographs showing double labelling for Iba1 (DAB, brown) and GS (Nickel-enhanced DAB, purple) in coronal sections, along the rostro-caudal axis of the IC, from animal RR4. The sub-regions of the IC shown overlaid (dashed lines) were drawn with reference to the atlas of Paxinos and colleagues (2009). (A) The rostral-most section shows strong GS neuropil labelling in the superficial layers of SC, with moderate labelling in the deeper layers and the BIC, and lighter labelling in the EC. Strong GS neuropil labelling was found in the PAG. (B-D) The outermost layers of DC and EC had strong GS+ neuropil labelling with less in CN throughout the rostro-caudal axis of IC. The leptomeninges can be seen to express selective Iba1+ labelling (B-D). Dorso-ventral and medio-lateral axes are shown in panel (C).

All three IC sub-regions were present in sections midway along the rostro-caudal axis (Figs. 1B-C). Stronger GS+ neuropil labelling was found in DC and EC than in CN. At the level where the ICs are joined by the commissure of the inferior colliculi (Fig. 1B) light bands of commissural fibres intercalated with GS+ neuropil banding along the medio-lateral axis. At this level, the lateral lemniscus had a similar pattern of labelling coursing along the dorso-ventral axis. Fibres of passage formed a clear border medially with the adjacent peri-aqueductal grey. This pattern was maintained at more caudal locations. At the caudal pole of IC the adjacent cerebellum also exhibited strong GS+ neuropil labelling in the molecular layer (Fig. 1D).

### Inverse somatic densities of GS+ and Iba1+ somata

Higher magnification images revealed specific somatic labelling of both epitopes. As expected, regularly spaced Iba1+ cells with pronounced ramifications tiled the parenchyma (Figure 2A-C). The territories over which Iba1+ ramifications extended were largely nonoverlapping, matching the classic morphology of sensing, non-activated microglia. Interspersed between Iba1+ ramifications were more numerous GS+ somata. Cellular GS+ labelling was constrained to the soma with a cytosolic distribution. In the overwhelming majority of cases, the nuclei of GS+ somata could be identified by an absence of staining, surrounded by dense cytosolic labelling.

**Figure 2.**
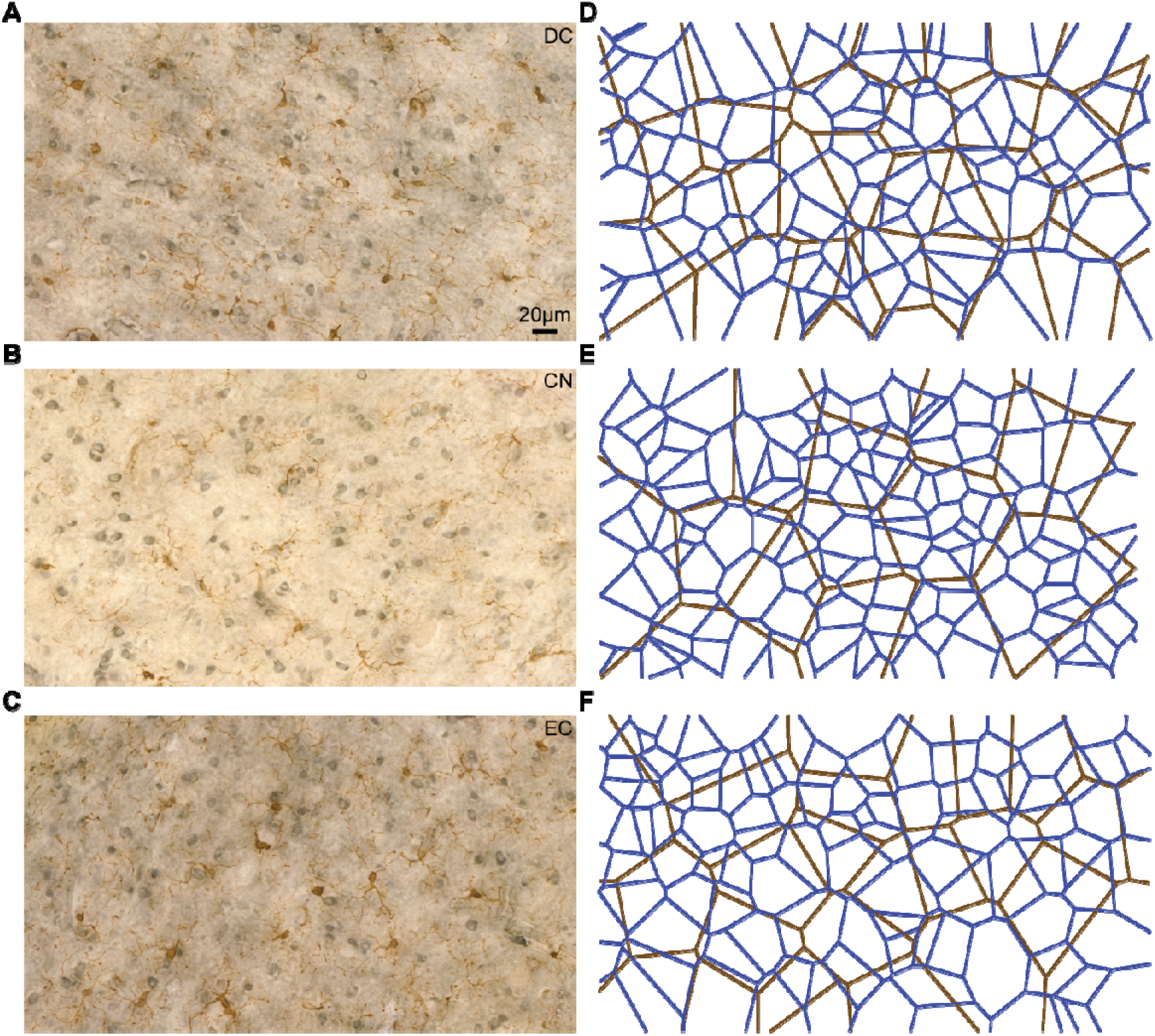
Photomicrographs (4Ox) showing representative examples of labelling in (A) DC, (B) CN and (C) EC. The density of Iba1+ cells was highest in DC, with the fewest in DC. Iba1+ cells tiled the parenchyma with ramified, non-overlapping processes, matching the known morphology of microglia. Conversely, GS+ cells had highest density in CN and lowest in DC. GS+ somata exhibited a non-uniform distribution, with examples of clustering as well as sparsely labelled regions. GS+ cells were more densely packed than Iba1+ in all regions. Note the lighter GS neuropil labelling in CN compared to the two cortical regions. (D-F) Voronoi tessellations of somata locations in left column for Iba1+ (brown lines) and GS+ (blue lines) somata. In all cases, Iba1+ Voronoi cells were homogeneous while GS+ territories were heterogeneous. Note the largest Voronoi cells for Iba1 and the smallest for GS were found in CN.

There was a clear difference in cellular density between both epitopes, with GS+ somata being more numerous. The DC consistently exhibited the greatest density of parenchymal Iba1+ somata (mean 64 somata per 100μm2; SD±9) and the lowest density of GS+ (83 ±15) (Fig. 2A). In contrast, the CN had the opposite trend, with the lowest density of Iba1+ somata (41 ±6) and the highest density of GS+ (119 ±17) (Fig. 2B). Cell densities in EC were nearer to the values found in CN than DC, with Iba1+ having a mean soma density of 48 (±5), with GS+ being 113 (±11) (Fig. 2C). Placement of soma location markers on regions of interest allowed derivation of Voronoi tessellations. Overlaying tessellations for each epitope across regions of interest (Fig. 2D-F) confirmed the greater density of GS+ somata than Iba1+ somata in all sub-regions, as well as the inverse relationship between both respective somata densities in DC and EC.

Soma densities of each cell type were semi-quantified by conducting manual cell counts from each IC sub-region of every section of all seven animals. A mean of 308 (±67) Iba1+ cells per IC were counted across 388 regions of interest. A total of 9,690 Iba1+ cells were counted with a minimum of 182 per IC sub-region. The mean number of Iba1+ somata per 100μm^2^ from each animal across IC sub-regions is shown in Figure 3A. The DC had the highest density of Iba1+ somata in all seven animals and the CN the lowest. A Welch’s one-way ANOVA with IC sub-region as the factor showed that the difference between these values likely reflected a real difference (W(2,11.6)=14.5; P=0.0007). Dunnett’s T3 multiple comparisons found these differences were between the DC and CN (t=5.5; adjusted P=0.0007) as well as DC and EC (t=4.1; P=0.006).

**Figure 3.**
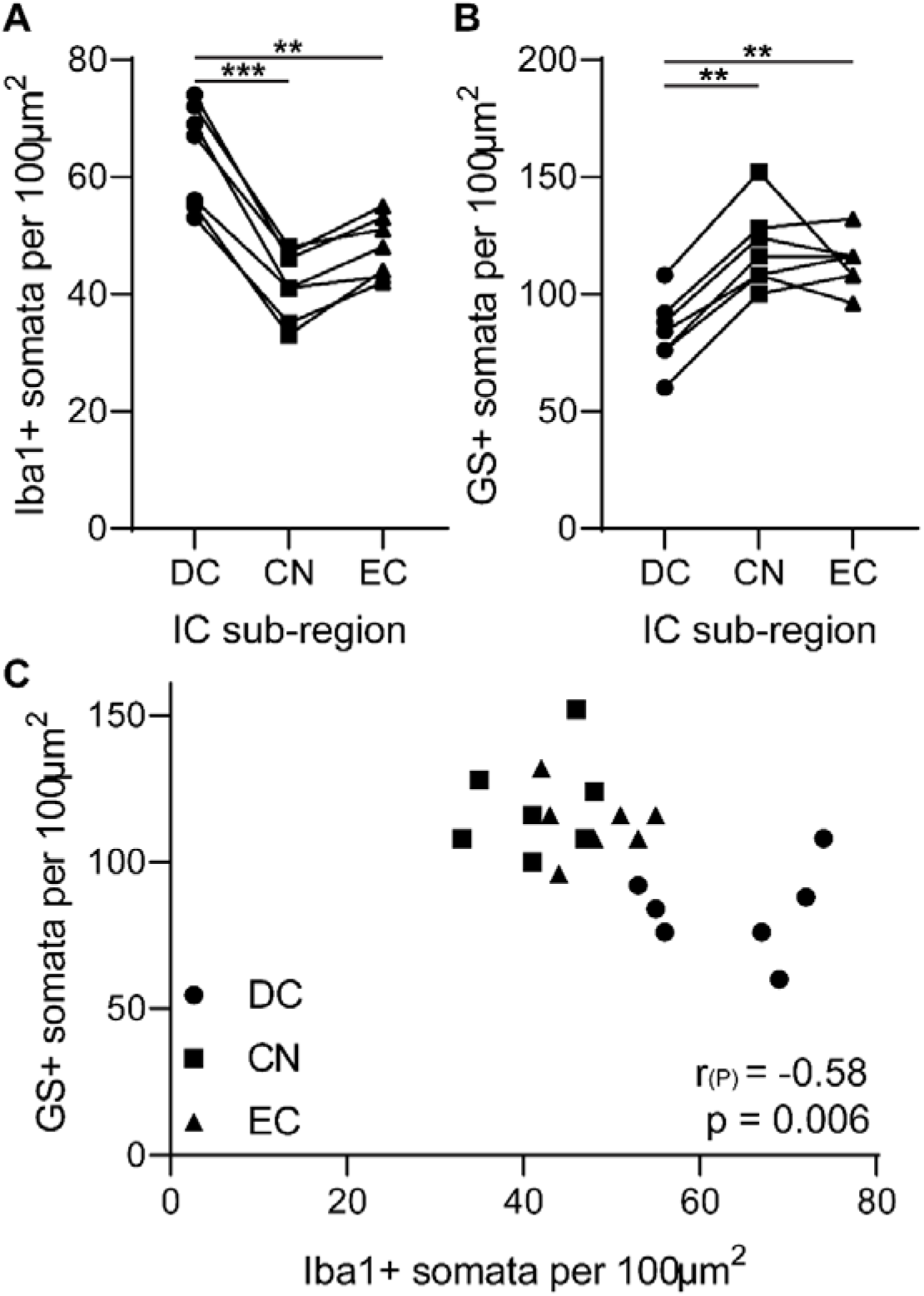
Inverse cell densities of Iba1+ and GS+ cells in IC sub-regions. (A) Iba1+ somata were more densely packed in DC than CN and EC for all seven animals. Mean values from each animal are plotted for each IC sub-region. Lines connect sub-regions from the same animal (n=7). (B) The reverse trend was found for GS+ cells, with the lowest density of soma counts in DC. (C) Plotting mean GS+ somata density as a function of mean Iba1+ somata density demonstrated a negative correlation between the two.

For GS+ cells a mean of 242 (±98) somata per IC across 194 regions of interest were counted. A total of 5,085 Iba1+ cells were counted with a minimum of 92 per IC sub-region. The mean number of GS+ somata per 100μm^2^ from each animal across IC sub-regions is shown in Figure 3B. The CN and EC had similar densities of GS+ somata in all seven animals. DC had the lowest GS+ somata density in all animals. A Welch’s one-way ANOVA with IC sub-region as the factor showed that the difference between these values likely reflected a real difference (W(2,11.5)=10.8; P=0.0023). Dunnett’s T3 multiple comparisons found these differences were between the DC and CN (t=4.1; P=0.004) as well as DC and EC (t=4.2; P=0.004).

Plotting mean GS+ somata count per 100μm^2^ as a function of Iba1+ somata counts demonstrated an inverse relationship in cell density between somata expressing the two epitopes (Fig. 3C). Pearson’s correlation revealed a negative relationship (r(P)=-0.58; P=0.006), due to DC having higher Iba1+ and lower GS+ counts and CN and EC having the opposite trend.

### Dorsal and external cortices have stronger neuropil labelling than central nucleus

To semi-quantify the variation in neuropil labelling, whole IC images were converted to 8-bit and the pixel grey levels along two axes on sections midway along the rostro-caudal extent of the IC were calculated. The first axis was placed dorso-ventrally through DC and CN along the classically defined low-to-high frequency sound axis in IC (Fig. 4A). The mean of three such lines through each IC are shown in a scatterplot in Figure 4B. The mean of all animals (black line) shows in initial decrease from the *glia limitans externa* to the outer border of the DC, indicative of strongest neuropil labelling 100-200μm into the parenchyma. This was followed by a monotonic, near linear increase in grey pixel values, demonstrating a gradual transition from DC to CN. Data from different animals are shown in different colours, with the same trend found in all cases. Pearson’s correlation revealed a positive association (r(P)=0.63; P=<0.0001). A second axis was placed orthogonally to the first and spanned from the lateral border of the IC to the medial border with the peri-aqueductal grey. A similar decrease was seen around 100μm from the *glia limitans externa*, followed by a near linear increase to midway along the axis. This was followed by a slight decrease and plateau along the medial half of this axis (Figure 4C). This difference between neuropil labelling along this axis through CN was characterised by a weaker correlation (r(P)=0.20; P=<0.0001).

**Figure 4.**
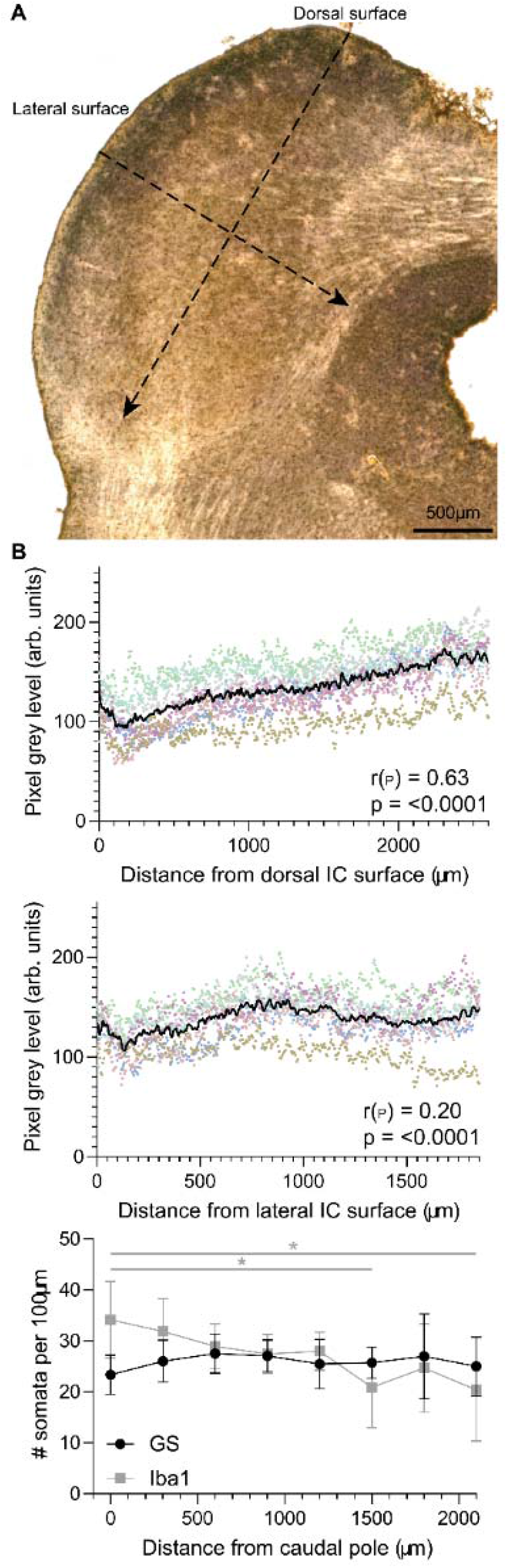
Neuropil labelling is greatest in the cortices and lightest in ventral CN. (A) Low power (5x) photomicrograph showing the IC labelling midway along the rostro-caudal axis from case RR2. Images were converted to 8-bit and pixel intensity calculated (5μm^2^ per pixel) along the dorso-ventral axis of the IC (dashed line), from the *glia limitans externa* to the ventral border of CN (arrowhead). (B) Neuropil labelling was darkest (lowest values) in the outer layers of the DC and thereafter followed a linear trend of decreasing intensity. Dots show individual measures. Each colour represents data from a different animal. All seven cases were included. Black line shows the mean of all seven cases. (C) The darkest labelling along the medio-lateral axis through IC was also found in the outer cortical layers. This was followed by a gradual decrease in labelling density to mid-way along this axis, followed by a plateau that extended to border with the *peri*-aqueductal grey. (D) Somatic density along the rostro-caudal axis of the IC was similar for GS+ cells but was significantly higher at the caudal pole for Iba1+ cells than in rostral regions.

Neuropil labelling also changed along the rostro-caudal axis, with stronger staining in DC and EC and weaker labelling in CN. Soma density counts were taken along the rostro-caudal axis at distances from 0 to 2,100μm from the caudal pole (distances calculated when the tissue was sectioned), as shown in Figure 4D. The density of GS+ somata were similar along the rostro-caudal axis but Iba1+ somata had highest density caudally and were lowest rostrally. A two-way ANOVA with cell type and distance from the caudal pole as factors found a two way interaction (F(7, 86)=2.3; P=0.03). Tukey’s multiple comparisons test revealed that the true effect was likely between Iba1+ somata counts at the caudal pole and those found at the rostral regions (0μm vs 1,500μm: q=5.3; P=0.023. 0μm vs 2,100μm: q=5.5; P=0.015).

### Peri-vascular somata density was similar across IC sub-regions

The above analyses considered cell counts in regions of interest where no vessels were present in the section. However, as both microglia and astrocytes are known to line blood vessels, a separate analysis was performed where longitudinal cross-sections through vessels were found in each IC sub-region, for all sections in all cases. The 2D distance between perivascular somata of GS+ and Iba1+ were respectively semi-quantified. Examples of longitudinal vessels were found throughout all sub-regions, with many extending over long distances (mean 2D length 211μm ±127). Vessel lengths were variable, with similar values in DC (198μm ±42); CN (241μm ±74) and EC (205μm ±18). Examples of typical vessels and adjacent perivascular labelling are shown in Figure 5(A-D). Despite the varied topology of IC sub-regions, the distance between both cell types was similar.

**Figure 5.**
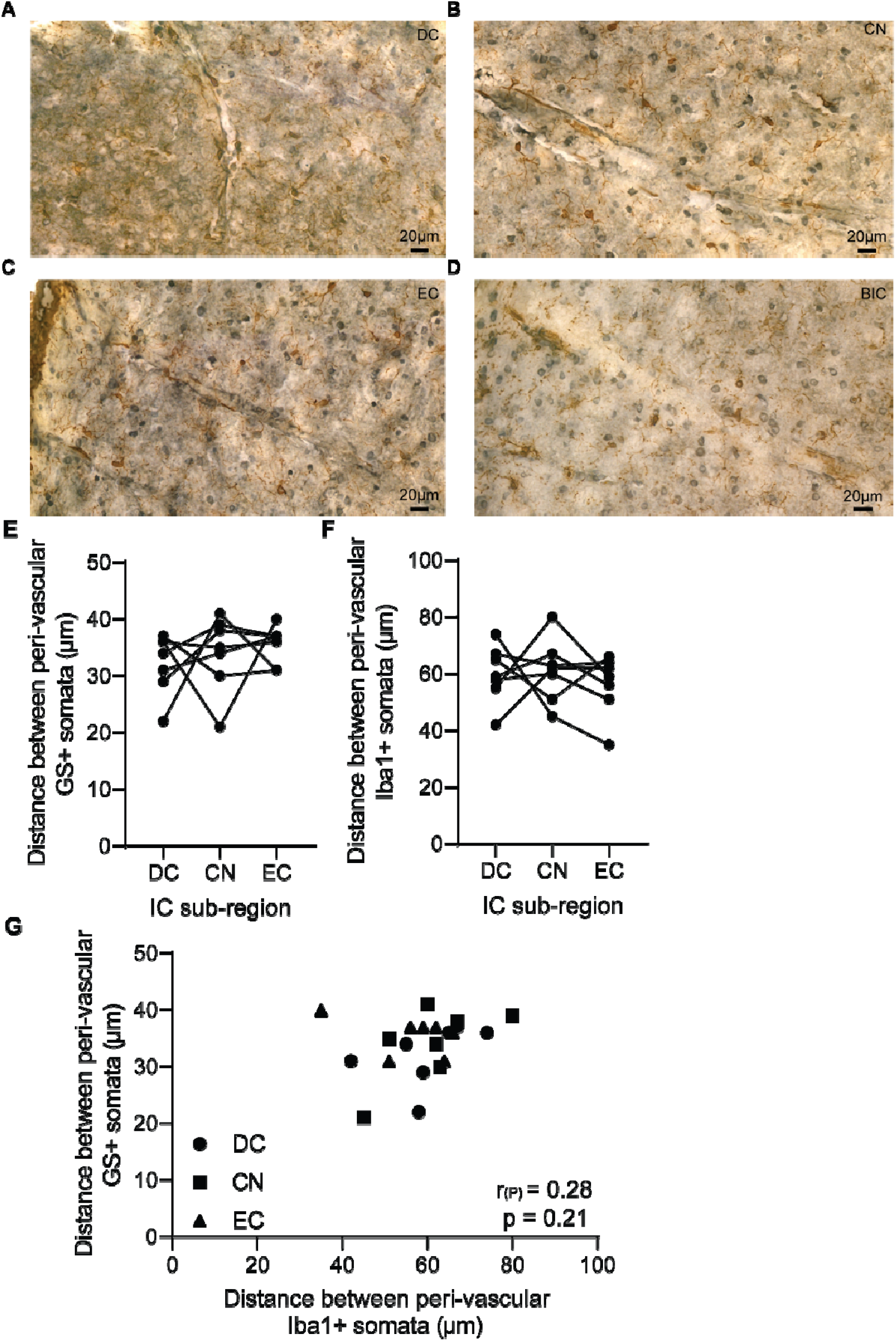
*peri*-vascular GS+ cells have around double the density of Iba1+ cells. (A) Representative example of *peri*-vascular labelling from DC and adjacent parenchyma. Note the presence of both Iba1+ and GS+ perivascular somata. Similar patterns of labelling were observed throughout IC, in (B) CN, (C) EC and (D) BIC. (E) 2D distance between peri-vascular GS+ somata revealed similar values across IC sub-regions, with a mean of 33.9μm. (F) There was also no difference between sub-regions for 2D distance between Iba1+ somata, but they were almost half as frequent, with a mean of 59.1μm. (G) Plotting 2D distance of GS+ somata as a function of Iba1+ somata revealed no correlation between the two.

The mean 2D distance between GS+ somata was 34μm (±16). This was similar in DC (32μm ±5), CN (34μm ±7) and EC (36μm ±3) (Fig. 5E). Welch’s ANOVA with sub-region as the factor demonstrated a lack of difference in peri-vascular GS+ soma density throughout IC (W(2, 11.0)= 1.0; P=0.40). As with parenchymal labelling, Iba1+ somata had half the density of GS+ somata. The mean distance between Iba1+ somata was 59μm (±23), which was similar in DC (60μm ±10), CN (61μm ±11) and EC (56μm ±11) (W(2, 12.0)=0.4; P=0.68) (Fig. 5F). A Pearson’s correlation also showed the lack of association between peri-vascular cell type density (r(P)=0.28; P=0.21).

## Discussion

These are the first descriptions of the distribution of GS+ somata and neuropil labelling in IC. Furthermore, building upon a recent report of Iba1+ microglia in guinea pig (Webb and Orton, 2020), these data reveal four novel features of putative glial labelling. Firstly, GS+ somata are more numerous than Iba1+ somata throughout the parenchyma. Secondly, there is an inverse relationship between GS+ and Iba1+ cellular density, with GS+ somata density greatest in CN, while Iba1+ density is greatest in DC. Thirdly, GS+ neuropil labelling is strongest in the cortices and weakest in CN with a gradual transition between sub-regions. Fourthly, the density of peri-vascular GS+ somata is double that of Iba1+ somata throughout IC. These data enhance our understanding of sub-regional variations in putative glial cell densities in IC.

### Inverse cellular density of astrocytes and microglia

The ubiquitous and fundamental role astrocytes and microglia play in the higher auditory pathway remains underexplored. Recent findings have revealed new roles for these glial sub-types in sensory nuclei. This includes evidence for astrocytes responding in a stimulus dependent manner in somatosensory cortex *in vivo* (Lines et al., 2020). Astrocytes have also recently been shown to modulate thalamic firing patterns *in vitro* via GABAergic signalling (Kwak et al., 2020). These findings coincide with advances in our understanding of the fundamental roles microglia play in sculpting synapses in the visual pathway in an activity dependent manner (Cheadle et al., 2020; Tremblay et al., 2010). In the auditory pathway, pharmacological depletion of microglia leads to deficits in early synaptic pruning at the calyx on Held (Milinkeviciute et al., 2019). However, further elaboration on the roles of astrocytes and microglia in higher auditory nuclei is lacking. An important challenge to understand the anatomical and physiological nature of the IC is the characterisation of glial cell types in the higher auditory pathway.

The present study suggests significant heterogeneity of astrocytes and microglia cell density between sub-regions of IC. Astrocyte heterogeneity takes many forms, including varied morphologies, developmental lineages, neurochemistry, and gene expression, as wells as the response to inflammation and neurological disease (Clarke et al., 2021; Miller, 2018). Similar degrees of heterogeneity are also found throughout the microglial population (Masuda et al., 2020; Silvin and Ginhoux, 2018). This poses a challenge to understand how glial sub-types contribute to developmental, ongoing and pathological processing in their local niche.

A surprising finding was the inverse somatic density between GS+ and Iba1+ somata. This suggests the functional roles of these cell types are antipodally related to each other across IC sub-regions. The high density of GS+ somata in CN could be due to an astrocytic role in refining and supporting auditory sensory processing. Certainly the highest proportion of GABAergic somata in IC, and perhaps anywhere in the brain, are located in ventral CN (Gleich et al., 2014; Ito et al., 2009; Webb and Orton, 2020). A greater number of GS+ astrocytes may be needed in CN to mediate the GABA shunt alternative to the citric acid cycle to support local GABAergic neurotransmission (Schousboe et al., 2013). Astrocytic control of biosynthesis and metabolism of GABA is well-established and a greater density of astrocytes in CN may be an adaptation to meet the demands of this local neural processing. Indeed SR101-expressing putative astrocytes in IC are known to express neurotransmitter transporters GAT-1 and GAT-3, which have key roles in GABAergic neurotransmission (Ghirardini et al., 2018), suggesting a role in GABAergic processing. However, astrocytes multiplex a range of physiological roles, including neurovascular coupling (Otsu et al., 2015; Petzold and Murthy, 2011). As the CN has amongst the density of capillaries (Song et al., 2011), the highest basal rate of oxidative metabolism in the brain (Sokoloff et al., 1977), and a high density of mitochondrial cytochrome oxidase expression (Cant and Benson, 2006, 2005). The high cellular density of putative GS+ astrocytes observed here may also be related to extensive neurovascular coupling in CN to meet ongoing high metabolic demand, which requires further exploration.

### Limitations and technical considerations

In anatomical studies such as this, the choice of target epitopes are of paramount importance to define the possible interpretation of observations. Antibodies for a number of distinct astrocyte epitopes are available, such as ALDH1L1, GFAP, GLAST, GLT-1, GS, NDRG2 and S100β. These markers label astrocytes with varying degrees of specificity and overlap with each other as well as other cell types.

Few parenchymal astrocytes in IC express GFAP, and those that do are located in the outer regions of DC and EC (Hafidi and Galifianakis, 2003; Webb and Orton, 2020). As somatic GS is known to predominantly label astrocytes (Martinez-Hernandez et al., 1977), with key roles in glutamate metabolism (Zhou et al., 2020), this marker was employed in the present study. However, GS is also expressed by some oligodendrocytes (Tansey et al., 1991; Xin et al., 2019), which may limit the strength of some interpretations of these data. However, the low density of GS labelling in the commissure and lateral lemniscus (Figure 1) suggests this may not be a major confounding factor.

There appears to be no marker that is entirely selective for IC astrocytes, which can currently circumvent this issue. Indeed, SR101 and S100β, which have also recently been used to identify putative astrocytes (Ghirardini et al., 2018) are also not entirely selective markers of astrocytes (Hill and Grutzendler, 2014; Steiner et al., 2007). This issue also applies to activated microglia, which express markers shared with macrophages, such as CD45 and CD11b. Consequently, a multiplexed approach that phenotypes glia based on multiple epitopes may be needed to better characterise sub-types of these cellular populations.

### Functional implications for neurochemical homeostasis

The gradual transition of GS+ neuropil labelling from around 100μm inside the parenchyma from the pial border of DC and EC, towards the CN (Figure 4) was also unexpected. In the distal outermost 100μm of IC, little GS+ neuropil labelling was observed, with a sharp border to the highest density of labelling proximally. The near linear reduction from this peak to lower GS+ neuropil in CN is in direct opposition to GS+ somatic density (Figure 3) and suggests two distinct pools of GS in IC - one somatic and another distributed throughout the neuropil.

It is striking that the highest density of GS+ neuropil is located in cortical regions. Cortical regions have few GABAergic neurons and a high density of NADPHergic and nNOSergic neurons (Coote and Rees, 2008). There is also a greater amount of glutamate (Adams and Wenthold, 1979) and NMDA receptors (Monaghan and Cotman, 1985) in DC than in ventral CN. Interestingly, the adjacent peri-aqueductal grey, an exclusively excitatory nucleus absent of GABAergic neurons, also exhibited strong GS+ neuropil, which formed a clear border with medial CN (Figs 1&4), highlighting neurochemical similarities with DC and EC. The different metabolic pathways required for local glutamatergic (putative excitatory) and GABAergic (putative inhibitory) neurotransmission suggest a role for the GS+ neuropil in glutamate-glutamine metabolism, as well as nitrergic and ammonia signalling and recycling. Nitric oxide (NO) has been shown to act as a volume transmitter at the calyx of Held (Steinert et al., 2008), which likely holds true for other regions of the auditory pathway, including the high density of nNOS+ neurons in the outer layers of DC and EC. Reactive oxygen species can inactivate GS (Schor, 1988) and NO is a small molecule whose actions are non-receptor specific and likely serves as a major redox source in DC and EC. Synaptic glutamate release into the extracellular space and consequent activation of NMDA receptors may lead to NO release from nNOSergic neurons (Olthof et al., 2019). This in turn may lead to oxidative stress and/or hyperammonemia in DC or EC (Suarez et al., 2002). The parenchyma is highly vulnerable to hyperammonemia and consequent oedema and perturbed ion buffering and GS plays a key role in nitrogen and ion homeostasis, alongside key roles in neurotransmission. Consequently, a high concentration of GS in synaptic, extracellular fluid or bound to the extracellular matrix may serve to buffer NO and maintain parenchymal nitrogen homeostasis. Indeed, modulation of GS by NO may directly modulate local glutamate metabolism (Kosenko et al., 2003), and in turn, excitatory neurotransmission in IC.

Another functional implication of GS+ variations between the DC/EC and CN are reports that 3-chloropropanediol-induced lesions lead to loss of astrocytes in CN but not the cortices of IC (Willis et al., 2004). The toxic effects of 3-chloropropanediol are thought to be due to disrupted redox signalling (Skamarauskas et al., 2007). The presence of high levels of GS in DC and EC neuropil may act to protect against such toxicity and may indicate a greater resilience of cortical IC regions to physiological redox stress than CN, with implications for plastic reorganisation of IC subsequent to hearing loss. This may include the greater proportion of descending input to DC and EC (Bajo et al., 2007). Corticofugal input to IC mediates plastic changes (Bajo et al., 2010) and is a candidate pathway to remediate some of the central effects of peripheral hearing loss (Lesicko and Llano, 2017), which may involve GS and nitrergic signalling.

## Contributions (CRediT)

LDO: Conceptualization, Data curation, Formal analysis, Funding acquisition, Methodology, Project administration, Resources, Supervision, Validation, Visualization, Writing - original draft, review and editing.

## Acknowledgements

The author thanks Dave Maskew, Megan Ryan and Rosie Mitchell for their technical support.

## Abbreviations

BIC: brachium of the inferior colliculus
CB: cerebellum
CoIC: commissure of the inferior colliculi
CN: central nucleus of the inferior colliculi
DC: dorsal cortex of the inferior colliculi
EC: external cortex of the inferior colliculi
GS: glutamate synthetase
Iba1: ionized calcium binding adaptor molecule 1
IC: inferior colliculi
LL: lateral lemniscus
PAG: peri-aqueductal grey
PBS: phosphate buffered saline
SC: superior colliculus

